# Single-cell metabolomics reveals the metabolic heterogeneity among microbial cells

**DOI:** 10.1101/2021.11.08.467686

**Authors:** Xuanlin Meng, Fei Tao, Ping Xu

## Abstract

In microbial research, the heterogeneity phenomenon is closely associated with microbial physiology in multiple dimensions. For now, A few studies were proposed in transcriptome and proteome analysis to discover the heterogeneity among single cells. However, microbial single cell metabolomics has not been possible yet. Herein, we developed a method, RespectM, based on discontinuous mass spectrometry imaging, which can detect more than 700 metabolites at a rate of 500 cells per hour. While ensuring the high throughput of RespectM, it integrates matrix sublimation, QC-based peak filtering, and batch correction strategies to improve accuracy. The results show that RespectM can distinguish single microbial cells from the blank matrix with an accuracy of 98.4%, depending on classification algorithms. Furthermore, to verify the accuracy of RespectM for distinguishing different single cells, we performed a classification test on *Chlamydomonas reinhardtii* single cells among allelic strains. The results showed an accuracy of 93.1%, which provides RespectM with enough confidence to perform microbial single cell metabolomics analysis. As we expected, untreated microbial cells will spontaneously undergo metabolic grouping coherence with genetic and biochemical similarities. Interestingly, the pseudo-time analysis also provided intuitive evidence on the metabolic dimension, indicating the cell grouping is based on microbial population heterogeneity. We believe that the RespectM can offer a powerful tool in the microbial study. Researchers can now directly analyze the changes in microbial metabolism at a single-cell level with high efficiency.

## Main

Heterogeneity widely exists in natural cells ^1, 2^. This phenomenon can be found in multiple aspects of genome ^3, 4^, transcriptome ^5, 6^, proteome ^7^, and metabolome ^8, 9^. In microbial research, heterogeneity is generally considered a crucial factor in the evolution and stability of cell populations ^10, 11^. Recent studies have extended the realization of heterogeneity effects in several dimensions, beyond being a critical factor in bioprocess robustness ^11, 12^, and cast light on the roles in drug resistance ^13^, quorum sensing ^14^, symbiosis ^10^ and adaptive evolution ^15^. Whereas cellular heterogeneity influences bioprocess performance in ways that to date are not entirely elucidated ^16^. This situation indicates the urgent demands for practical single cell tools that can easily explore cellular heterogeneity.

However, there are few single-cell microbial research methods compared to mammals, which restrains the intuitive access to the essence of microbial heterogeneity. Significantly, the lack of protein and metabolite single cell methods derivates the obstacle of direct PPI (protein-protein interaction) network and MRN (metabolite reaction network) reconstruction to counteract the cognition of cellular heterogeneity. In the past decade, some works have been conducted applying Matrix Associated Laser Desorption Ionization (MALDI) mass spectrometry to characterize metabolites at a single cell level ^8, 9, 17-19^. However, few studies were performed to address the defects of the poor repeatability among chemical matrix background and the immaturity of analytical solution for microbial single cell data analysis ^20^, which are the primary obstacles of establishing microbial single cell metabolomic method.

Herein, we constructed a new method, “RespectM,” to perform single cell metabolomics research in microorganisms. In our RecpectM method, single cell data collection is precisely achieved by discontinuing mass spectrometry imaging (MSI). A sublimation method was applied for MSI preprocessing to deal with the problem of poor matrix repeatability. To establish the analytical system, standard software SCiLS Lab (Bruker, Germany) and open accessed R packages were integrated to form the pipeline ^21-23^. These mature software and packages endorsed our method standardization and more readability in the data processing. To promote the accuracy of data acquisition, we used microscopy-guided cell ablation coupled with in situ micron-level precise positioning.

Recently, researchers have artificially constructed a microbial population, and further analysis under energy-constrained conditions found that the cell population can respond to energy limitation in a predictable manner, which likely contributes to the stability and robustness of microbial life ^24^. Inspired by this work, we conducted a single cell metabolomic analysis of the *Chlamydomonas reinhardtii* population to explore microbial metabolic heterogeneity. Herein, we established different *Chlamydomonas reinhardtii* cell groups in three dimensions: temperature, cell wall-deficient, and photosynthesis diversity within the total cell populations. RespectM provided a data set of 4302 cells and more than 700 metabolites. All metabolite identifiers are chosen following RespectM standards, which contain lipids and metabolites with KEGG annotations. The identified series of features include neutral glycerolipids (DG, TG), protective lipids (PE, CerP), signal transduction lipids (PI, PIP), nucleotides (GMP), pigments (Porphyrin), and metabolites belonging to the central metabolic pathway (Oxaloacetate). The above results indicate that the metabolite identification ability of RepectM can reach the level of bulk metabolomics. Finally, 111 features were selected from dysregulated pathways for downstream single cell analysis.

Significantly, the results show that RespectM can distinguish single microbial cells from the blank matrix with an accuracy between 95.3% to 98.4%. This result shows that using RespectM method can avoid matrix interference in the single cell data acquisition process. We applied the cubic support vector machine (SVM) method to classify two allelic *Chlamydomonas reinhardtii* mutants at a single cell level ^25^. In allelic strains, we can achieve 93.1% identification accuracy.

Further, we applied a UMAP chart to display the basal metabolic status of single cells and introduced pseudo-time analysis to find the cell trajectory ^26-28^. Ultimately, we captured the cell trajectory and the dysregulated metabolite accumulation induced cell grouping (DMACG) phenomena by stream algorithm. Based on UMAP analysis, we obtained information about the accumulation of metabolites in specific single cells. Capturing these DMACG can provide us with new knowledge for exploring the heterogeneity among microbial cells. In future microbial research, researchers can design precise metabolite synthesis for each microbial single-cell metabolic profile.

## Results

### The RespectM method

The main challenges of MALDI-based single cell metabolomics include precise and complete single cell data acquisition, repeatable micron-scale matrix applications, and enough peaks for identification. Herein, we composed laser etching guided droplet microarray (LEM), DHB matrix sublimation, and sparse data matrix generation to the RespectM method to meet with above challenges (Fig.1a(i)) ^29^. Although studies attempted to do precise positioning ^30, 31^, however, more precise positioning is demanded when MSI is applied to micron-scale microorganisms. Therefore, the traditional marking using drawing and mechanical stress marking needs to be improved. Herein, the laser was applied to array a pattern on the surface of ITO glass slides. Then we use fine needles to copy the pattern after DHB matrix application to obtain micron level positioning. We can perform simple cell liquid dripping in the designated area according to the pattern’s guidance. Another crucial technical detail is to calculate the laser offset. Since single cell data acquisition uses a laser raster of <50 μm for data acquisition, laser should be bombarded in the middle of the cell to avoid incomplete single cell acquisition. All in situ images are collected under the stereomicroscope.

**Fig. 1.**
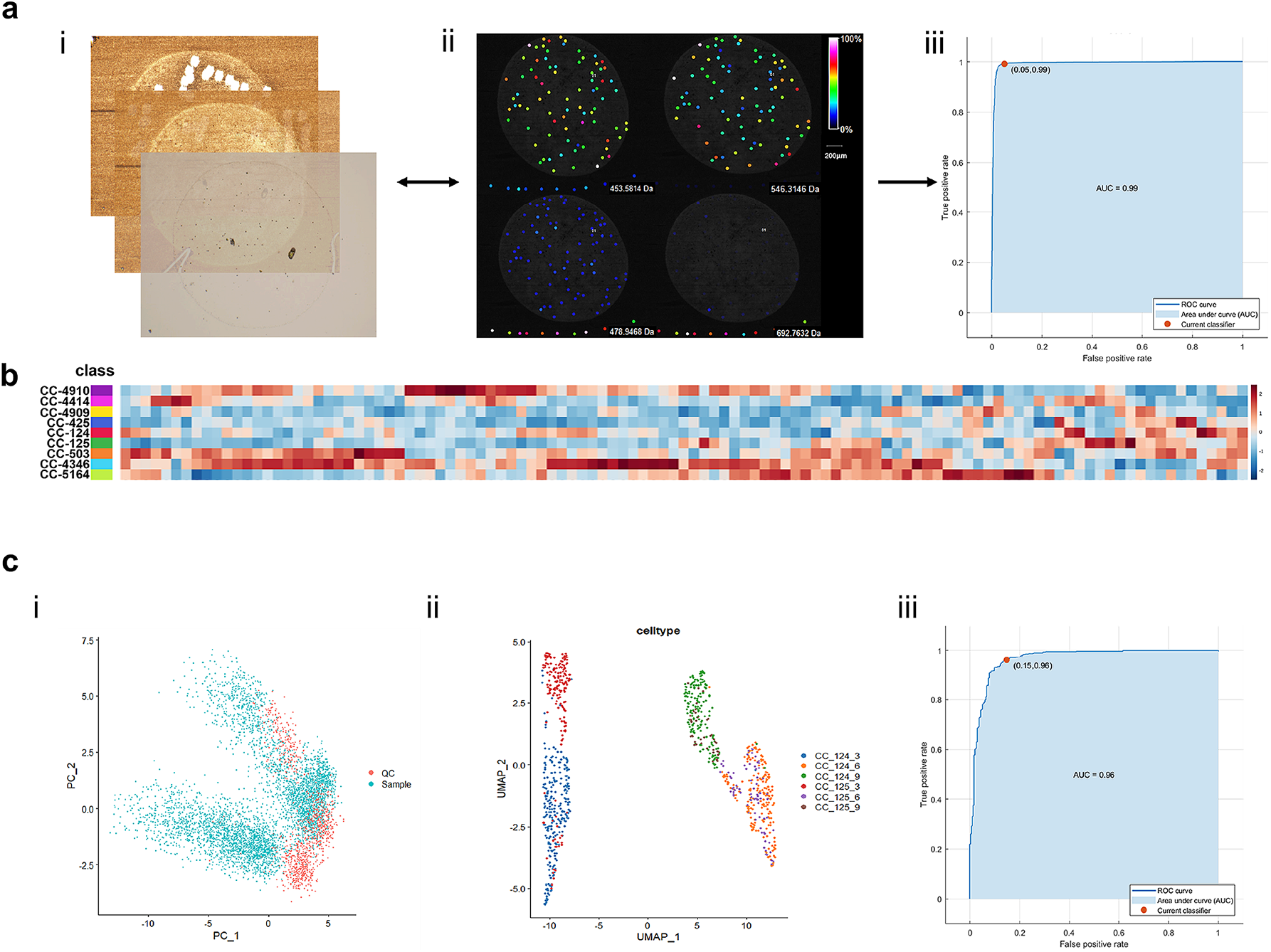
The workflow and validation of RespectM method. RespectM integrates a high-resolution stereomicroscope and MALDI-MRMS for discontinuous MSI acquisition to obtain in situ metabolic spectra of every single microbial cell. The QC-based correction strategy can provide a unique single-cell metabolic profile despite matrix effects. (i) Correspondence between the original optical image, the optical image after sublimation, and the optical image after MSI data collection. (ii) Visualization of matrix-cell relationship. It includes the amount of two cells>matrix and two cells<matrix feature in terms of abundances. Scale bar, 200 μm. (iii) Matrix-cell classification test based on Cubic SVM. **b**, Annotation of dysregulated features based on LC-MS/MS. **c**, RespectM can distinguish two *Chlamydomonas reinhardtii* with one allelic differenc**e**. (i) PCA mapping between matrix and sample. (ii) UMAP visualization of CC-124 and CC-125 in three-time points. (iii) RespectM can perform single-cell allelic difference classification accuracy up to 93.1%.

Studies have shown that continuous MSI strategies can be performed efficiently in the in situ data acquisition ^32-34^. However, cross-contamination between adjacent raster will reduce the confidence of MSI data. The small size of microbial cells aggravates this phenomenon. Therefore, we use the discontinuous MSI acquisition function of fleximaging (Bruker, Germany). This mode can automatically collect batches of in-situ single cell data from non-adjacent pixels and proceed by SCiLS Lab (Bruker, Germany) software in a short time (Fig.1a(ii)). In this study, we applied discontinuous MSI to collect *Chlamydomonas reinhardtii* single cell metabolic data under the premise of laser offset calibration and micron-level positioning. Besides, ten blank matrix points were collected as QC samples at the end of each dripped spot sequence to avoid the batch effect of discontinuous MSI. It is also the first time we proposed QC-based MALDI metabolomics (Fig.1a(ii)). Finally, RespectM can provide a single cell data matrix corresponding to in situ MSI acquisition sequence. The reads include the sequence of data acquisition, in situ position of the single cell in the corresponding series, and the peak annotation based on MS and MS/MS identification results (Extended Data.1).

### RespectM can distinguish allelic strains among single cells

In MSI research, the chemical matrix is necessary to associate the desorption of biological samples, but it also complicated the background in the MSI data. It is worth noting that many studies have neglected the signal interference of chemical matrix covered on biological samples. In this research, we propose a new solution to retain the features of S/Q>1.2 in the sparse matrix. It can be reputed that the feature signal response in the sample is higher than the feature in the blank chemical matrix.

Further, the sample data needs to be distinguished from the blank matrix to ensure the accuracy of downstream single cell analysis. One hundred eleven metabolites were chosen to correspond to LC-MS/MS metabolomic and lipidomic for classification (Fig.1b). On this basis, 22 classification strategies were applied to distinguish samples from the blank matrix. The results show that these methods can achieve 95.3%–98.4% classification accuracy (Fig.1a(iii)). Furthermore, RespectM was tested for the ability to distinguish allelic strains CC-124 and CC-125 (Fig.3c(i-ii)). Through the cubic support vector machine (SVM) strategy, 93.1% classification accuracy of two strains can be achieved (Fig.3c(iii)).

Further, we conducted Seurat to perform analysis on CC-124 and CC-125. Since the reproduction time of *Chlamydomonas reinhardtii* is 24 hours, the mixed cell state at the three time points will inevitably undergo significant changes. According to the instruction of Seurat, the principal component analysis (PCA) reduction should be advanced before uniform manifold approximation and projection for dimension reduction (UMAP). The UMAP chart shows that the distance of single cell data between three time points is significant. Then we visualized the marker features used for automatic grouping. We applied the Wilcoxson algorithm to calculate all the 16 markers in auto-clusters. Results have shown that the amount of these markers were dysregulated accumulated in the auto-clusters. We found that the relative intensity of marker features is consistent under the premise of wild-type *Chlamydomonas reinhardtii* without external stimulation. In parallel, some parts of cells clustered respectively in each time point, indicating the variability of single cell metabolic adaptation.

### Use RespectM to visualize the heterogeneity in cell populations

The metabolic heterogeneity of microorganisms lacks an effective visualization method. This situation also limits the characterization of the metabolic status of different microbial populations at the single cell level. Taking *Chlamydomonas reinhardtii* as an example, the metabolism of cells within the population is highly variable, which elaborated the collection of the single cell metabolic status. Besides, the cells sampled at different time points are in a mixed state.

We use RespectM to visualize the cell population based on UMAP to cope with this obstacle. The result has shown that there was no apparent clustering of single cells at the three and nine day (Fig.2a (i, iii)). However, two clusters occurred in the cell population at the six day (Fig.2a (ii)). Combined with species analysis, it was found that the three different groups of *Chlamydomonas reinhardtii* had significant metabolic changes at the six day (Fig.2a (ii)). In the temperature group (CC-124, CC-4414, CC-5164), there is no apparent clustering of single cells at the three and nine day. It is worth noting that two clusters occurred in the six day (Fig.2b (ii)). One group comprises three *Chlamydomonas reinhardtii*, and another is the cell group CC-4414 and CC-5164. Here we focused on the joint clusters CC-4414 and CC-5164, where the accumulation of metabolites helps to explain the reasons for clustering. Therefore, we visualized the markers used to distinguish different clusters in the entire cell population of the temperature cell group.

**Fig. 2.**
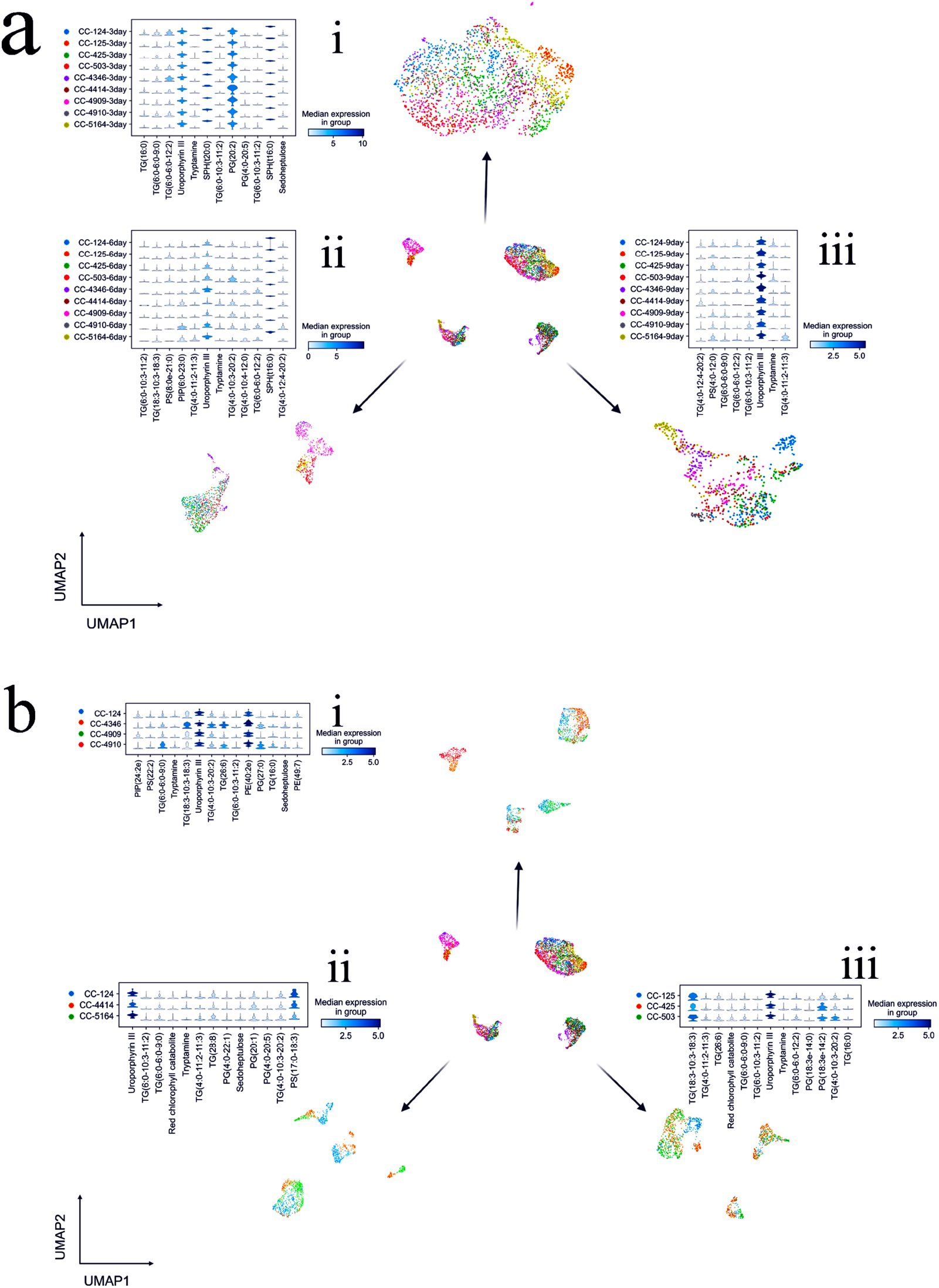
Single cell metabolism heterogeneity of *Chlamydomonas reinhardtii* among time points and species. **a**, Visualize the heterogeneity of the total cell population at three-time points. (i) Markers and cell population at the three day. (ii) Markers and cell population at the six day. (ii) Markers and cell population at the nine day. **b**, Visualize the heterogeneity of the total cell population at three groups. (i) Markers and cell population at the photosynthetic group. (ii) Markers and cell population at temperature group. (ii) Markers and cell population at the cell-deficiency group.

Further, by visualizing the TOP 50 high variable features, we found that the three lipids of DG (8:1e-11:1), PE (36:0e), PS (22:3-13:0) gradually decreased with time. The change of PG (20:1) is on the contrary. It is worth noting that CerP (d16:1-25:0) is a unique marker in two cell groups at the six day; In the photosynthetic group (CC-124, CC-4909, CC-4910, CC-4346), we found 14 marker features combined with high variables for analysis and visualization (Fig.2b(i)). We focused on the two cell groups generated in the six day. One group consists of CC-4909 and CC-124, and the other group is CC-4910 and CC-4346. The two groups of the six day are independent, respectively (Fig.2b(ii)). In the distribution of metabolite abundance at the six day, we found that PIP (24:2e) has high abundance in CC-124 and CC-4909 cell group, while PE (16:1-12:4) is highly accumulated in CC-4910 and CC-4909. CerP (d16:1-25:0) is mainly accumulated in the two cell groups at the six day. As time goes by, the abundance of DG (8:1e-11:1), PG (16:0-18:3), PS (22:3-13:0) are decreased, while PG (20:1) on the contrary; In the final cell wall defect population, the 6-day cell population did not generate a cluster obviously (Fig.2b(iii)).

Nevertheless, consistent with the temperature and photosynthetic groups, the abundance of PG (20:1) accumulated with time and is on the contrary of DG (8:1e-11:1). In this section, we visualized the metabolic heterogeneity of *Chlamydomonas reinhardtii* at the single cell level. However, UMAP analysis cannot reveal the detailed microbial cell trajectory.

### Identification of *Chlamydomonas reinhardtii* DMACG phenomenon by RespectM

RespectM provides a more intuitive representation of the heterogeneity of single cell metabolism. The above part mainly shows the heterogeneity of *Chlamydomonas reinhardtii* in the two dimensions of time point and category. However, due to the complex cell mixing state at each time point, more heterogeneity information is hidden in the cell population (Fig.3a). Due to the rapid reproduction of microorganisms, the metabolic state is unstable compared to higher organisms. This phenomenon complicated the capture of the metabolic state. Therefore, we have applied a matured pseudo-time method to explore further microbial heterogeneity in the cell trajectory (Fig.3b, c, d).

**Fig. 3.**
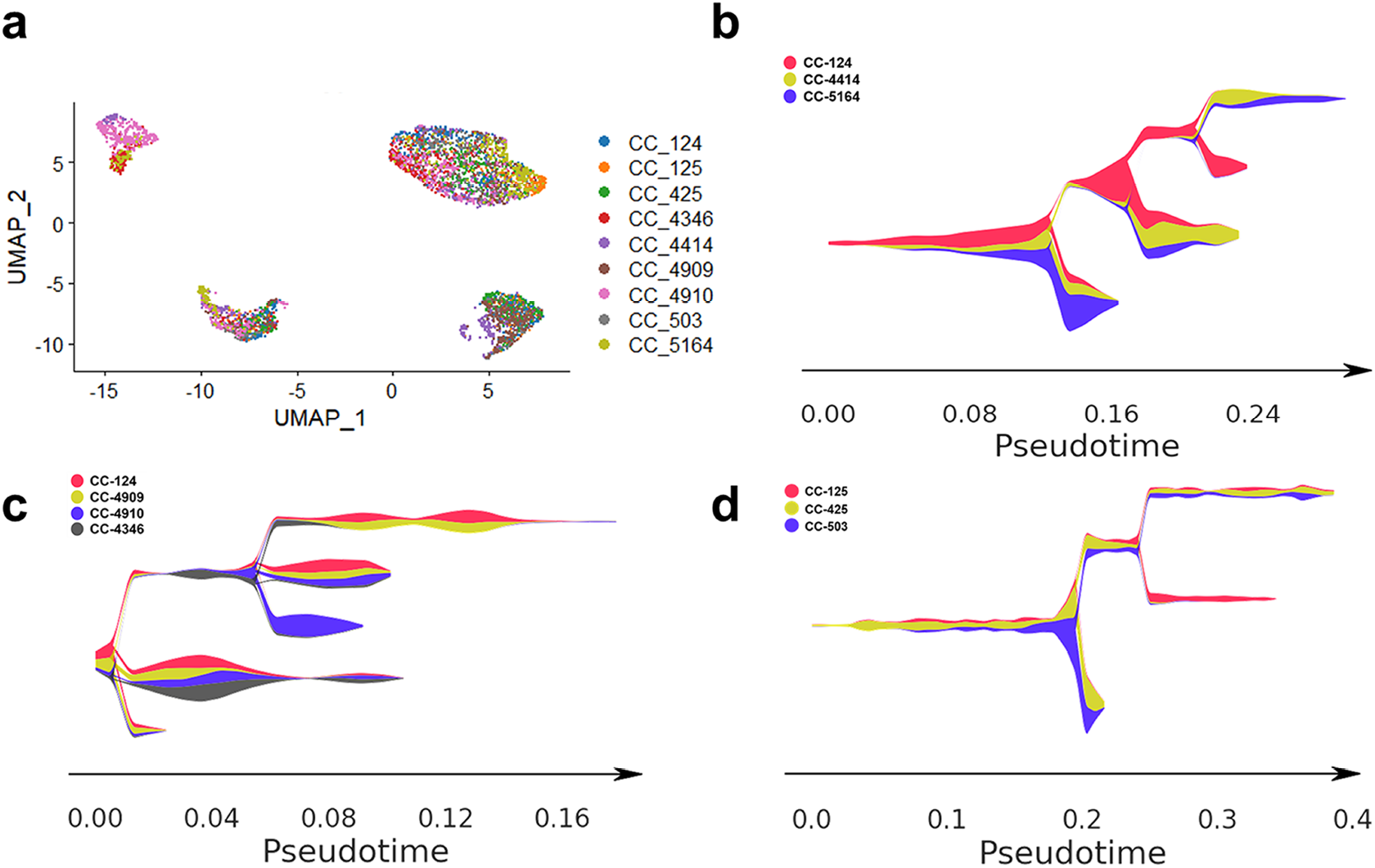
RespectM reveals DMACG phenomenon among *Chlamydomonas reinhardtii* single cell population. **a**, Visualization of 4302 *Chlamydomonas reinhardtii* single cells based on 111 metabolites. **b**, Temperature group contains three groups of *Chlamydomonas reinhardtii*: CC-124, CC-4414 and CC-5164. Pseudo-time analysis reveals the different branches of normal temperature and very temperature adapted strains. **c**, Photosynthesis group contains four groups of *Chlamydomonas reinhardtii*: CC-124, CC-4909, CC-4910 and CC-4346. Pseudo-time analysis reveals the branches among the three lines which have a coherent genetic relationship. **d**, Cell wall-deficiency group contains three groups of *Chlamydomonas reinhardtii*: CC-125, CC-425 and CC-503. Pseudo-time analysis reveals the different branches of wild type and cell wall-deficiency strains.

CC-124, CC-4414, and CC-5164 at the S5 node have different DMACG phenomena in the temperature group (Fig.3b). CC-124 independently generated an S5-S7 branch, while CC-4414 and CC-5164 form an S5-S6 branch. The results show that the intensity of TG, DG, PC, PG in the S5-S6 branch is higher than that of S5-S7, and the abundance of PE, TG, PIP, CerP, LPC in the S5-S7 branch is higher than that of S5-S6 in parallel. This phenomenon shows that CC-4414 and CC-5164 produce more PE under normal temperature conditions to adapt to the environment. Meanwhile, the high abundance of phytic acid and PIP lipids confirms a difference in phosphatidylinositol metabolism between CC-124, CC-4414, and CC-5164 at the single cell level. It is worth noting that LPC, the precursor of PC, accumulated in the S5-S6 branch in high abundance. In contrast, PC has a high abundance in S5-S7, which implies that the high & low-temperature group adapted to normal temperature metabolism through the limitation of LPC conversion.

In the photosynthetic group, the inheritance of CC-124, CC-4909, and CC-4910 is continuous. This relation is reflected in the S0-S2 and S3-S5 branches (Fig.3c). It is worth noting that CC-4909 and CC-124 shared the same branch, which is consistent with the information that CC-124 is the background strain of CC-4909. However, the CC-4909 based mutant strain CC-4910 has a unique branch S3-S6 beside three joint branches with CC-4909, which is different from the S3-S4 branch and has dysregulated abundance in LPC, PE, PG, PS, TG, PC. This phenomenon is in line with the description that CC-4910 has a stronger metabolic capacity. It is mainly shown in lipid synthesis, transportation and storage capacity, and stronger metabolic capacity than the other two strains. In parallel, CC-4346 shares the same branch with the other three Chlamydomonas at S0-S1. However, S0-S1 shows a higher lipid abundance than S3-S5 branches overall. This phenomenon implies that CAO deficiency demands CC-4346 to enhance metabolism to make up for its photosynthetic defects.

CC-425 and CC-503 are wall-deficient types belonging to the S3-S4 branch (Fig.3d); The cells of wild type CC-124 (with cell walls) are mainly distributed in the S0-S1 branch. The abundance of N-Carbamoyl-L-asparate in the wall-deficient type CC-425 and CC-503 is higher than that of the wild type of CC-124, which means that the wall-deficient type needs stronger primary metabolism to resist the defect of the wall. In the S3-S4 branch, the abundance of TG, DG, PE, PC are higher than S0-S1, but CerP, PG, LPC are the opposite. In general, lipids are a vital part of the cell membrane, so that the LPC component is converted to the PC component in the cell wall-deficient strains to enhance the membrane fluidity^35^. It is worth noting that when the nine *Chlamydomonas reinhardtii* are analyzed in a pseudo-time series without comparison, the basal status of *Chlamydomonas reinhardtii* still produces the DMACG phenomenon in the cell trajectory. This result means that we can apply RespectM method to reveal the basal metabolic heterogeneity by reordering single cells.

## Discussion

To explore the heterogeneity of cells, researchers have tried some top-down & bottom-up strategies in mammals ^36-38^. However, the lack of microbial single cell metabolic fingerprint restrains the visualization of microbial metabolic heterogeneity. This work aims to energize microbial single cell metabolomic research by developing the RespectM method. The advantages of RespectM include several dimensions. For instance, the LEM cell chip strategy simplified the preprocessing steps before MSI data collection and optimized the single cell in situ positioning accuracy during MSI data collection. Further, A confidence grading system including the MS and MS/MS identification strategies were developed for single cell metabolic identification. Moreover, the QC-based MSI calibration method in MALDI-based single cell metabolomics was proposed. Altogether, all these factors above gifted RespectM unique advantages in convenience, accuracy, and throughput. By establishing the RespectM methodology, we can identify more than 700 metabolites in more than 500 single cells within one hour.

In single-cell MSI data acquisition, the cleanliness of the matrix background and accurate cell positioning is the basis. For now, the spray coating strategy is the common matrix application method for microbial single cell research ^9, 30^, which is not applicable for microbial cells. Bien et al. conducted a systematic evaluation of the existing single-cell matrix application strategies. The results showed that the matrix application of the sublimation strategy could best retain the in-situ and precise metabolic information ^20^. Therefore, RespectM chose the matrix sublimation method. This method has the weakest matrix delocalization effect and the best signal repeatability in MSI. Compared with the newly proposed strategy “spaceM”, RespectM has re-established the microbial single-cell metabolomics method and has unique advantages in the matrix application section.

At present, some factors are restricting the application of MSI in microbial single-cell research. For instance, the size of a single microbial cell is only 1-10 μm, while the raster ablation size is usually around 50 μm. Therefore, optical figure splicing operation is inappropriate to microbial single cell data acquisition. Taking Bruker ultrafleXtreme as an example, the physical movement accuracy of the mechanical stage is 5 μm; optical image stitching will also bring micron-level errors. This information indicates that while we expand MSI in situ range, positioning errors increases as well.

To ensure the cleanliness of single cell data, we first compared RespectM with methods requiring cell staining and fixation ^30^. RespectM has a shorter preprocessing process, and the cell sample is obtained through regular shake flask cultivation. Both cell bulk metabolomics and single cell metabolomics are directly sampled from culturing cells at the same time. Secondly, we applied the discontinuous MSI method in RespectM to prevent cross-contamination between adjacent data points in data collection. The accuracy distinguishing single cell from blank matrix reaches 98.4%; this result offers RespectM enough confidence in microbial single cell acquisition. Thirdly, there is no method for batch correction between MSI internal data points. Therefore, MALDI-based single cell data collection needs to cope with a new batch correction strategy. We introduced matrix background points as QC to correct batch effects between multiple data acquisitions. This strategy differs from the batch correction method of single cell transcription, and mature metabolomics algorithms MetNormalizer gifted RespectM with higher confidence.

After verifying the accuracy of the methodology, we used RespectM to perform single cell metabolic analysis on the artificially constructed three cell populations of *Chlamydomonas reinhardtii*. The cellular heterogeneity is visualized on the two dimensions of sampling time points and different cell groups. First, all single cells generated 4 clusters on the UMAP graph (Fig.3a). There is no apparent grouping between the three and nine day (Fig.2a(i, iii)). It is worth noting that there was a clear grouping in the six day (Fig.2a(ii)). This result implies that the six day should be the most prominent point in time when heterogeneity occurs. We further analyzed the endogenous heterogeneity of the three artificially constructed cell populations, respectively. Interestingly, the results showed that the number of cell clusters has increased (Fig.2b). Therefore, we propose that the cognition of heterogeneity will change by the observation angel.

Moreover, all the *Chlamydomonas reinhardtii* microbial single cells were reordered through pseudo-time analysis. Compared with mammal metabolic pseudo-time analysis, the metabolic pseudo-time analysis is more diverse. The results showed a reordered DMACG phenomenon consistent with the biochemical and genetic relationships among three cell populations. Without external stimulation, the metabolic heterogeneity of *Chlamydomonas reinhardtii* produced DMACG in different dimensions. The dysregulated accumulated metabolites in different single cells, combined with the “cell age” gifted by pseudo-time analysis, allows us to analyze, for example, the phenomenon of joint clustering of single cells from different species and obtain new knowledge.

In all, RespectM not only fills the large yet empty niche in the chaotic modulating stage of microbial single cell metabolomics but has already helped in characterizing the DMACG phenomenon in *Chlamydomonas reinhardtii*. Beyond the results validated in line with previous reports, a new dimension of knowledge that RespectM provided will flourish microbial research. The simple procedure, high compatibility with microbiology practices combined with an integrated bioinformatic pipeline will drive the democracy of single cell microbial metabolomics forward to a new era.

## Materials and Methods

### *Chlamydomonas reinhardtii* resources and growth conditions

*Chlamydomonas reinhardtii* was obtained from Chlamydomonas Resource Center University of Minnesota (http://www.chlamycollection.org) and Freshwater Algae Culture Collection at the Institute of Hydrobiology (FACHB) (http://algae.ihb.ac.cn/). All species of algae were grown up on a solid TAP mediμm, then transferred single clone to a liquid TAP mediμm for three replicates, respectively. Cells were cultured in a shaker under ∼100 μEinstein m ^-2^ s ^-2^, 100 rpm.

### Cell handling and microarray preparation

For microarray preparation, the preprocessing part of this method is quite simple. We designed patterns on the surface of the ITO slide and used lasers for pattern-based etching. As shown in the figure, Since the endogenous single cell status is different at each time point, we choose the three, six, and nine day for cell harvesting. While sampling for LC-MS/MS metabolomic and lipidomic, a small amount of cell fluid was taken to centrifuge and wash with deionized water. 10 mL *Chlamydomonas reinhardtii* cell culture was taken from the three, six, and nine day. Samples were centrifuged at 2000 x g for 5 mins and resuspended two times in deionized water to reduce the remaining TAP mediμm among cells. Then dilute cells to an appropriate density. After dripping 0.2-0.5ul cell suspension to the designated area of LEM, we placed the chips on the clean bench and waited 15-20 mins for the blow-dry. Before matrix sublimation, it is necessary to take optical graphs by stereomicroscope (Olympus & MVX10, Japan) of each cell droplet unit on the prefabricated cell chip. The purpose of this step is to recognize single cells in optical images.

### Matrix application and cell localization

After preparing microarrays, in this study, we use 2,5-Dihydroxybenzoic acid (DHB) as a matrix because it is suitable to analyze small molecules. Matrix was applied by sublimation on LEM. The parameter of sublimation was 181°C, 12mins (iMLayer, SHIMADZU, Japan). Since high-purity DHB is unstable in the air, we completed the downstream preparations within 20 minutes (https://www.aladdin-e.com/zh_cn/d119198.html).

To localize single cell on microarrays, we use a stereomicroscope to take optical images. Needles were applied to mark the original laser-etched pattern on the LEM surface. Hence the micron level positioning was achieved. Meanwhile, the calculation of laser spot offset is the prerequisite to collect single cell data automatically. Generally, the laser does not bombard at the absolute central in flexcontrol (Bruker, Germany), demanding that we calculate the relative offset from the experiment strike position to the absolute central position. After calculation, we can mark the cells on the adjusted positions in discontinue MSI acquisition mode, fleximaging (Bruker, Germany). Optical images were taken after matrix application for MSI in situ data acquisition. Further, single cells were recognized combined with original optical images.

### Single cell data acquisition by MALDI-MRMS

Throughout the experiment, MALDI-MRMS (SolariX 7.0T, Bruker) was used for high-resolution MSI acquisition. The constant temperature in the mass spectrometer room is 18°C, and the hμmidity is maintained between 50% and 60%. The instrμment parameter settings are as follows. Funnel RF Amplitude: 150 V; Time of Flight=0.9 ms. The laser offset calculation should be performed in advance before the experiment at different times. Then mark the acquisition point in the correct position on the fleximaging software. After data acquisition, optical images were taken to compare before laser ablation to ensure the accuracy of laser bombardment position. We collect ten blank matrix spots in each sequence for quality control. In parallel, 20 standards were selected due to their biological function and m/z. For instance, the m/z of A, B, C, D, E were 100, 300, 500, 700, 900. The selection of 5 m/z can ensure the confidence of calibration between 60-1500 m/z.

### Sample preparation

For the extraction of metabolites, we use 80% methanol as the solvent, sonicate and centrifuge at 3000 rpm for 15 minutes, and then take out the supernatant for centrifugation at 12000 rpm for 30 minutes; We use Folch reagent (chloroform: methanol=2:1) for lipid extraction then freeze-dried and re-dissolved in a solvent (chloroform: methanol=1:1).

### LC-MS/MS methods for metabolomics and lipidomics

This study applied Thermo UPLC Q-Extractive plus (QE) coupled with ESI ion source for ed metabolomics data acquisition. RP Zorbax Eclipse XBD-C18 colμmn was chosen for positive and negative mode analysis. In the mobile phase selection, we refer to the method of Luo et al., use 0.1% formic acid in diluted water as aqueous phase A and 0.1% formic acid in pure acetonitrile as organic phase B. The LC gradient elution program was as follows: t = 0.0 min, 99% A; t = 5.0 min, 99% A; t = 5.5 min, 70% A; t = 9 min, 100% B; t = 11 min, 100% B; and t = 12.1 min, 99% A.

The MS parameters of C18–ESIMS in the analysis of positive and negative ionization mode are as follows: the mass range was set from m/z 80 to 1000, with a spectra collection rate of 2.0 Hz and capillary voltage of 4500 V.

QE plus with ESI ion source was applied for ed metabolomics data acquisition as well. Agilent Poroshell EC-C18 colμmn (3 × 50 mm, 2.7 μm) was chosen for positive and negative mode analysis. Both the MS acquisitions of metabolomics and lipidomics were performed on Xcalibur (ThermoFisher, USA). The following parameters were set to correspond to Meng et al.

### The RespectM pipeline of single cell metabolite-lipid identification

Compound Discover and Lipidsearch commercial software were applied to identify metabolites and lipids, respectively. Identification results can be found in supporting information. In brief, we have classified the identification results of MALDI sc-metabolomics with different confidence levels. The metabolites and lipids that bulk omics can directly annotate are classified into Level 1 and 2, and the MS1 peaks identified by the metID algorithm are classified as Level 3 according to adduct annotation.

## Data analysis

For bulks omics analysis, MetaboAnalyst was chosen to do statistical analysis. In parallel, pathway analysis was performed within different cell groups. Details can be found in supporting information. For single cell matrix analysis, the RespectM mainly includes several parts. We imputed the sparse matrix by sclmpute algorithm, and applied QC sample to do batch effects correction. After matrix normalization, Seurat and Stream are used for downstream analysis.

## Author contributions

Xuanlin Meng performed the experiments and analyzed the data. Xuanlin Meng wrote the manuscript. Fei Tao and Ping Xu critically reviewed the manuscript and revised it. All authors read and approved the submitted version.

## Acknowledgements

This work was supported by the grants of National Key R&D Program of China (2018YFA0903600) and National Natural Science Foundation of China (22138007, 31870088 and 32170105).

